# Diminution of feeding response in *Helicoverpa armigera* by inhibition or silencing of sugar gustatory receptor

**DOI:** 10.1101/599324

**Authors:** Aniruddha R. Agnihotri, Sanyami S. Zunjarrao, Mukta Nagare, Rakesh S. Joshi

**Affiliations:** Institute of Bioinformatics and Biotechnology, Savitribai Phule Pune University, Pune, MS, India; School of Veterinary and Life Sciences, Western Australian State Agricultural Biotechnology Centre (SABC), Murdoch University, Perth, WA, Australia

**Keywords:** Gustatory receptor, *Helicoverpa armigera*, Miglitol, Feeding response

## Abstract

Gustatory receptor (GR) is one of the essential chemosensory molecules in Lepidopteran pests. GR is involved in sensing several canonical tastes which in turn regulate the diverse behavioral and physiological responses of these insects. In this article, we have evaluated the alteration in feeding response of *Helicoverpa armigera* by blocking and silencing of sugar-sensing gustatory receptor 9 (HarmGr9). Sf9 cells based assay showed that glucose analogue, Miglitol, can bind to the expressed HarmGr9. This binding might lead to an inhibition of receptor activation and downstream signaling, indicated by reduced intracellular Ca^2+^ fluorescence. Further, the in-vivo study illustrated the feeding rate reduced on a diet containing miglitol as compared to the larvae fed on the artificial diet. Reduction in feeding rate was prolonged in insects fed on the miglitol containing diet even after switching to the control glucose diet. Competitive cell-based assay and feeding assay, using equimolar glucose and miglitol, also showed an inhibitory effect on HarmGr-9 activation and insect feeding rate. We have observed similar feeding rate reduction in HarmGr9 knockdown in *H. armigera* larvae. We believe this unique approach of *H. armigera* feeding response inhibition by blocking the sugar receptor can be further used to develop a novel strategy for agricultural pest management.

## 1. INTRODUCTION

Lepidoptera is a significant insect order from an agricultural perspective. Their ecological success majorly depends on adaptive chemoreception tactics that play an essential role in the selection of hosts, egg-laying sites, and mates (Dahanukar et al., 2005). Some of the insects from this class exhibit a wide range of hosts and diverse feeding behavior, due to diverse and or highly specialized chemosensory mechanism.

Chemosensory system act as an interface between the insect and its environment as it stimulates signaling cascade in response to external chemical cues. Insects show two primary types of sensilla, the main sensory organ of insects, uniporous gustatory sensilla and the multiporous olfactory sensilla (Agnihotri et al., 2016). Gustatory sensilla are involved in eliciting various behaviors, which are broadly distributed on the mouthparts, antennae, wings, and ovipositor of insects (Dahanukar et al., 2005; Dethier, 1976; Scott et al., 2001; Stocker, 1994). The gustatory sensilla comprises of four GR neurons and one mechano-sensory neuron, which are involved in sensing the taste and other related stimuli (Agnihotri et al., 2016). Taste neurons have GRs on their dendritic end, which can fundamentally differentiate between the host plant nutrients and the deterrents. Different type GRs are responsible for sensing various canonical tastes like bitter, sweet, salty, sour, and umami. Sweet taste detecting GRs are involved in identifying the food sources rich in sugar and this further leads to attractive or positive feeding responses in insects (Sato et al., 2011; Xu et al., 2012). It has been observed that multiple sweet GR can be single sugar specific and conversely multiple sugars can be detected by single GR. (Dahanukar et al., 2007; Jiang et al., 2015; Jiao et al., 2008; Slone et al., 2007; Wanner and Robertson, 2008).

In insects, Gr43a clades represent major sugar receptor class in the larval stage. These receptors are primarily characterized for sensing the fructose molecules. They are also present in the brain tissues to detect the fructose level in hemolymph and regulate feeding behavior of the larval insects (Miyamoto et al., 2012; Sato et al., 2011). Multiple GR candidates from this class have been identified in *Bombyx mori* (BmGr9) and *Helicoverpa armigera* (HarmGr4 and 9). Although these receptors belong to Gr43a family, they can respond to a wide range of non-fructose sugars (Freeman et al., 2014; Mang et al., 2016). It has been reported that HarmGr9 is involved in detecting multiple monosaccharides and disaccharides such as D-glucose, D-fructose, D-galactose, D-maltose and, sucrose (Xu et al., 2012).

Polyphagous insect pest, *H. armigera* (Hübner) (Lepidoptera: Noctuidae), is distributed worldwide and lead to average crop loss of approximately 2 billion USD annually (Jiang et al., 2015; Sharma et al., 2005; Xu et al., 2012). Due to remarkable chemosensory adaptation, these insects have developed enormous dietary flexibility (Agnihotri et al., 2016). The emergence of pesticide resistance and broadening of host range in *H. armigera* imposes challenges to agriculture productivity and highlights the need for effective controlling strategy.

In this report, we have assessed the feeding behavior of *H. armigera* upon blocking by iminosugar, Miglitol and siRNA mediated silencing of the sugar-sensing HarmGR9. We have performed the cell-based assay to validate the binding of miglitol in presence of glucose. Furthermore, we observed behavioral and physiological changes upon HarmGr9 blocker feeding. These results were then supported using RNAi technology, to understand the effect of blocking this sugar receptor on insect feeding ability. We believe this report would provide insights into *H. armigera* feeding behavior changes due to sugar receptor blocking and can be further explored to develop a novel additive strategy for agricultural pest management.

## 2. MATERIALS AND METHODS

### 2.1. Material

Miglitol (≥99.9 %, Medical grade) and Glucose (≥98 %) were obtained from Sigma Chemical Co., St. Louis, MO, USA. Analytical grade chemicals and culture media were purchased from HiMedia Laboratories, Mumbai, India. Molecular biology grade chemicals and enzymes were procured from Thermo Fischer Scientific, Massachusetts, United States.

*H. armigera* eggs were ordered from National Bureau of Agricultural Insect Resources, Bengaluru, Karnataka, India, and were reared on artificial diet. Laboratory conditions were maintained at humidity 60%, temperature 28 °C and photoperiod of 16 hours light: 8 hours dark for one generation as described in previous reports (Joshi et al., 2014).

*Spodoptera frugiperda* (Sf9) insect cell line was purchased from the National Centre from Cell Sciences (NCCS) cell repository and was maintained in Grace’s insect medium (HiMedia,) supplemented with 10 % inactivated fetal bovine serum (HiMedia) at 28 °C as a semi-adherent culture.

### 2.2. Isolation and Cloning of HarmGr9

*HarmGR9* gene cloned in PIB/V5-His (PIB/V5_Gr9) vector was obtained from CSIRO, Australia. The detailed methodology of gene isolation and cloning is described previously (Xu et al., 2012).

### 2.3. Calcium imaging assay

The method for calcium imaging was adapted from a previously reported article (Zhang et al., 2011). Sf9 cells were seeded in 12-well plate and were incubated for 48 hours at 28 °C. After the incubation period, the cells were transfected with 700 ng of plasmid construct (PIB/V5_Gr9) and 3 μl of FuGENE^®^ HD transfection reagent (Promega, USA) in 100 μl of medium per well. After 48 hours of incubation, post-transfection, calcium imaging and data analysis were performed as previously described (Zhang et al., 2011). Fluo-4 AM (Thermo Fischer Scientific) and DAPI dihydrochloride (HiMedia) were the fluorescent dyes used to carry out calcium imaging and nuclear staining respectively. Glucose (natural ligand) was used as a positive control, and miglitol (Glucose analogue, which can compete with glucose for binding on HarmGr9) was used a test ligand. Both the ligands were used at 50 mM concentration.

### 2.4. *In-vivo* feeding assay

Initially, all the mixed-sex fifth instar test and control larvae were starved for 8 hours. Post starvation miglitol and glucose were independently (each 50 mM concentration) fed to the larvae in 1 cm^2^ block of 1% agarose. The initial weight of the block and larvae was recorded followed by taking the weight of block and larvae for 12 hours, at the time interval of every 1 hour.

Competitive feeding assay was performed with miglitol and glucose (50 mM) together in 1 cm^2^ block of 1% agarose using the same methodology as mentioned above. The control insects for this experiment were fed on agarose block containing only glucose (50 mM).

### 2.5. Leaf disc feeding assay

For leaf disc feeding assay, fully grown fresh tomato plant leaves cut in disc shape of 1 cm of diameter were used. Leaf discs were first washed with sterile DEPC water and dried. Insects were starved for 8 hours and then were provided with one leaf disc per insect for feeding. Test insects were fed on leaf discs dipped and in 50 mM of Miglitol solution, whereas for feeding control insects, the leaf discs dipped in sterile DEPC water were used. Feeding rate and insects weight was recorded every 2 hours, for 8 hours.

### 2.6. RNAi studies and qRT-PCR analysis

HarmGr9-siRNA was synthesized by using MEGAscript™ RNAi Kit (Thermo Fisher Scientific, USA) following the manufacturer’s instruction. Mixed-sex fourth instar larvae were fed with HarmGr9-siRNA, 16μg per insect, in artificial diet, whereas DEPC water was fed to the control larvae. The insects were observed for 24 hours. Post this total RNA was extracted from the treated and control insects using SV Total RNA Isolation kit (Promega, USA). The RNA was quantified and qualified by a NanoDrop ND-2000 (Thermo Fisher Scientific, USA). The cDNA was synthesized using a RevertAid First Strand cDNA Synthesis kit (Thermo Fisher Scientific, USA) according to the manufacturer’s manual.

Relative transcript abundance of HarmGr9 in control and Gr9 silenced insects was determined by quantitative Real-Time PCR (qRT-PCR) using 7900HT Fast Real-Time PCR System (Applied Biosystems, Foster, CA, USA) and GoTaq^®^ qPCR Master Mix (Promega, GmbH, Germany). The relative expression of HarmGr9 was assessed (Joshi et al., 2014). B-actin (Accession No.: AF286059) was used as a reference gene for normalization. Reaction mix for qRT-PCR and thermal cycler conditions were followed as described in a previous article (Chikate et al., 2013). The average transcript abundance and subsequent fold difference with respect to the control were calculated.

### 2.7. Statistical Analysis

All the experiments were performed in two set of triplicates. Student’s t-test was used for statistical analysis. Data was expressed as mean ± SD. A p-value <0.05 was considered as statistically significant.

## 3. RESULT AND DISCUSSION

### 3.1. *In-vitro* blocking of HarmGR9 receptor activation

*In-vitro* expression of HarmGr9 in Sf9 cells by transfection followed with calcium imaging was carried out to study the alterations in downstream signaling pattern of HarmGr9 in the presence of natural and test ligands. Non-transfected Sf9 cells were used to compare the effect of test and control ligands on HarmGr9. Transfected Sf9 cells showed heavy calcium influx in the presence of glucose as compared to the non-transfected Sf9 cells (Fig. 1A). Also, when exposed to glucose, the transfected cells demonstrated the high intensity of fluorescence. Whereas, the intensity of fluorescence decreased significantly in transfected cells, when exposed to miglitol, thus supporting our hypothesis of miglitol being a Gr9 blocker (Fig 1A and B). This difference in calcium influx was also evident in total Relative Fluorescence Count (RFU) of transfected cells when exposed to glucose and miglitol separately (Fig 1B). Further, similar to the previous reports, background fluorescence in non-transfected cells in the presence of glucose due to the endogenous expression of taste receptors in Sf9 cells was observed (Xu et al., 2012).

**Figure 1:**
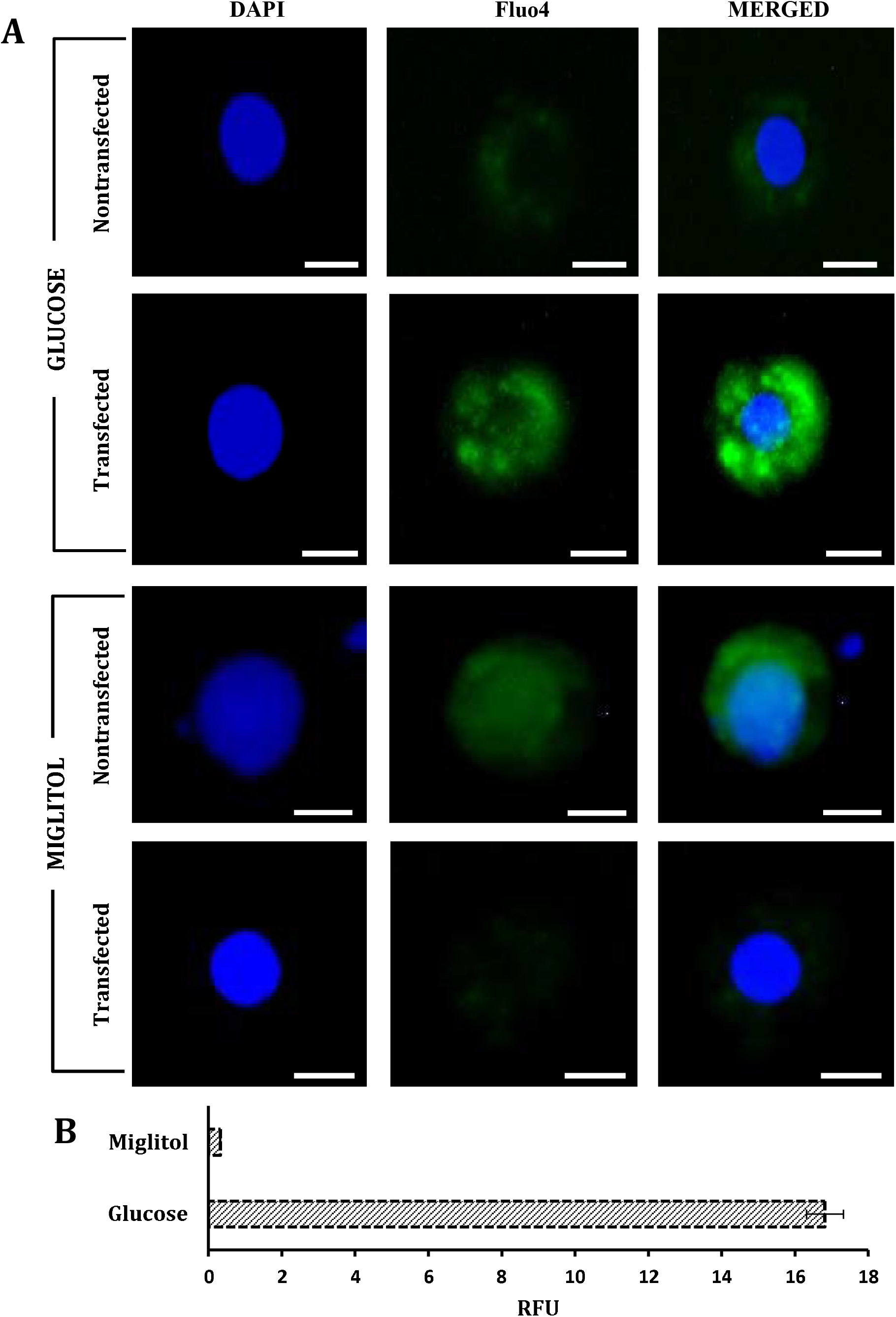
*In-vitro* blocking of HarmGr9 expressed in Sf9 cells. **(A)** Calcium imaging of transfected and non-transfected cells showing the difference calcium influx (green fluorescence) in presence of glucose and miglitol; Blue fluorescence indicate nuclease stained with DAPI; Both the ligands used in 50 mM of concentration individually. Scale bar =10 μm. **(B)** Change in calcium influx of transfected Sf9 cells in presence of glucose and miglitol represented in the form of Relative Fluorescence Unit (RFU)

The decrease in calcium influx of transfected cells in the presence of miglitol indicates that this ligand is a potential blocker of HarmGr9 and thus inhibits the activation of the downstream signaling cascade of this receptor. Comparatively, very little or no fluorescence was observed in both the non-transfected as well as transfected cells in absence of any treatment. A similar study was performed by Y Bobkov and E Corey in 2015 using selected insect odorant receptors/odorant co-receptors (Or/Orco) for the antagonist mediated blocking. KB-R973, a Na^+^/Ca^2+^ ion gated channel inhibitor was reported as an odorant evoked calcium influx inhibitor by showing the Or/Orco complex blocking in *Anopheles gambiae* (Bobkov et al., 2014). Our result illustrated the similar properties of miglitol as a HarmGr9 blocker and calcium influx inhibitor.

### 3.2. Miglitol negatively modulate feeding response of *H. armigera*

The fifth instar mixed-sex larvae of *H. armigera* were chosen for carrying out insect feeding assay and were starved for 8 hours before the experiment. Insects were fed on 1% agarose block (1×1 cm) containing single ligand, Miglitol (50mM) for test and glucose (50mM) for control. Insects showed voracious feeding of agarose block containing Glucose which is the natural ligand of HarmGr9. A significant drop of 50 – 60%, in the insect’s feeding rate, was observed when they were allowed to feed on agarose block containing Miglitol (Fig. 2A). Very little or no feeding was observed when the insects were fed only on agarose block. Reduction in feeding rate of insects in the presence of miglitol suggests that the blocking of HarmGr9 suppresses the chemosensory signals required for feeding and thus, leads to a decrease in their feeding potential.

**Figure 2:**
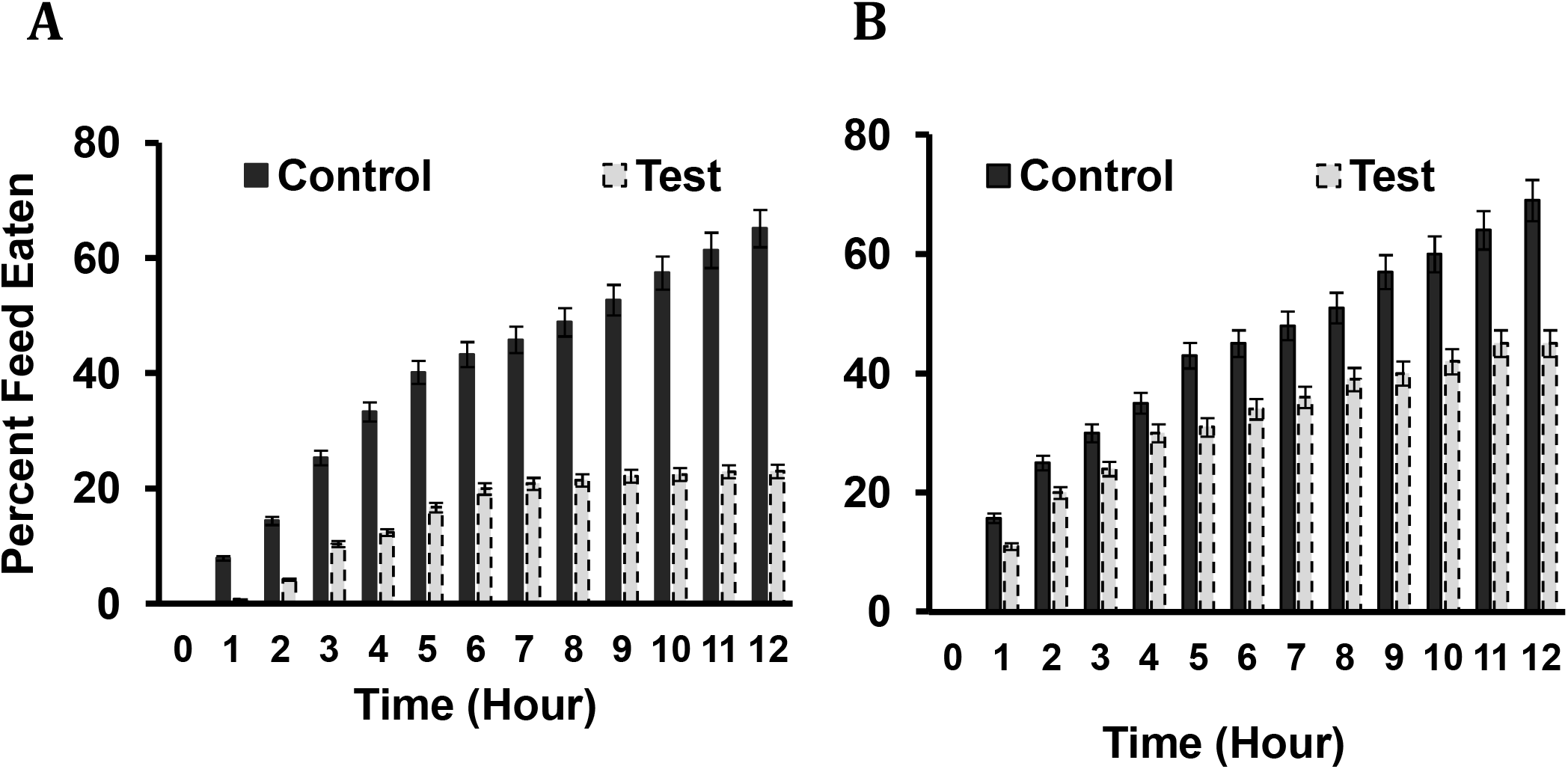
*In-vivo* effect of Miglitol on insect feeding response. **(A)** Feeding assay using agarose block containing glucose (control) and Miglitol (test); Both ligands used in 50 mM concentration individually. **(B)** Percent feed eaten by insects fed on agarose block containing glucose +miglitol (50 mM each) (test) and only glucose (control)

Further, competitive feeding assay was performed in order to check whether the decrease in feeding rate is due to the blocking of glucose specific receptor. In this, the insects were fed on agarose block containing glucose and miglitol together (both in 50 mM concentration). Simultaneous feeding of sugar and miglitol, sugar analogue, resulted in the similar blocking effect (as observed in the above result) of HarmGr9 in the insects. We observed a significant difference in feeding rate of test insects as compared to the control insects. In both the experiments we observed that initially, the difference in feeding rate of test and control insects is comparatively less (<5 %) followed with a significant increase in this difference in later hours. This difference gradually increased from 5% to more than 40% as the feeding continued up to 12 hours (Fig. 2B). According to us, this particular pattern was due to the localization of Gr9 in the foregut of the larvae (Xu et al., 2012). It took some window period for the blocker to reach the actual site of the receptor expression and till that period insect showed regular feeding in accordance with the control insects. Once the threshold number of receptors is blocked, the insect showed a gradual reduction in feeding rate, which remains almost constant throughout the experiments (Fig. 2A and B).

### 3.3. Effect of HarmGr9 blocking on host plant feeding

Tomato is one of the most targeted crop plants by *H. armigera*. Thus, in order to check the effect HarmGr9 blocking on the feeding response of these insects in the presence of the tomato plant, two experimental setups were designed. In first set up, the larvae were fed on fresh tomato leaf discs dipped in miglitol (test) and DEPC water (control). The feeding response was judged based on percent weight gain (PWG) of the insects during feeding. We observed a significant difference in the PWG of the test and control insects. The overall PWG by the test insects was around 35-40% less as compared to the control insects (Fig. 3A). We intentionally gave natural leaves as a feed to the insects, to check the inhibitory effect of HarmGr9 blocking in the presence of natural leaf sugars. The leaves were cut in a disc shape of 1 cm diameter to provide an equal amount of food to every insect.

**Figure 3:**
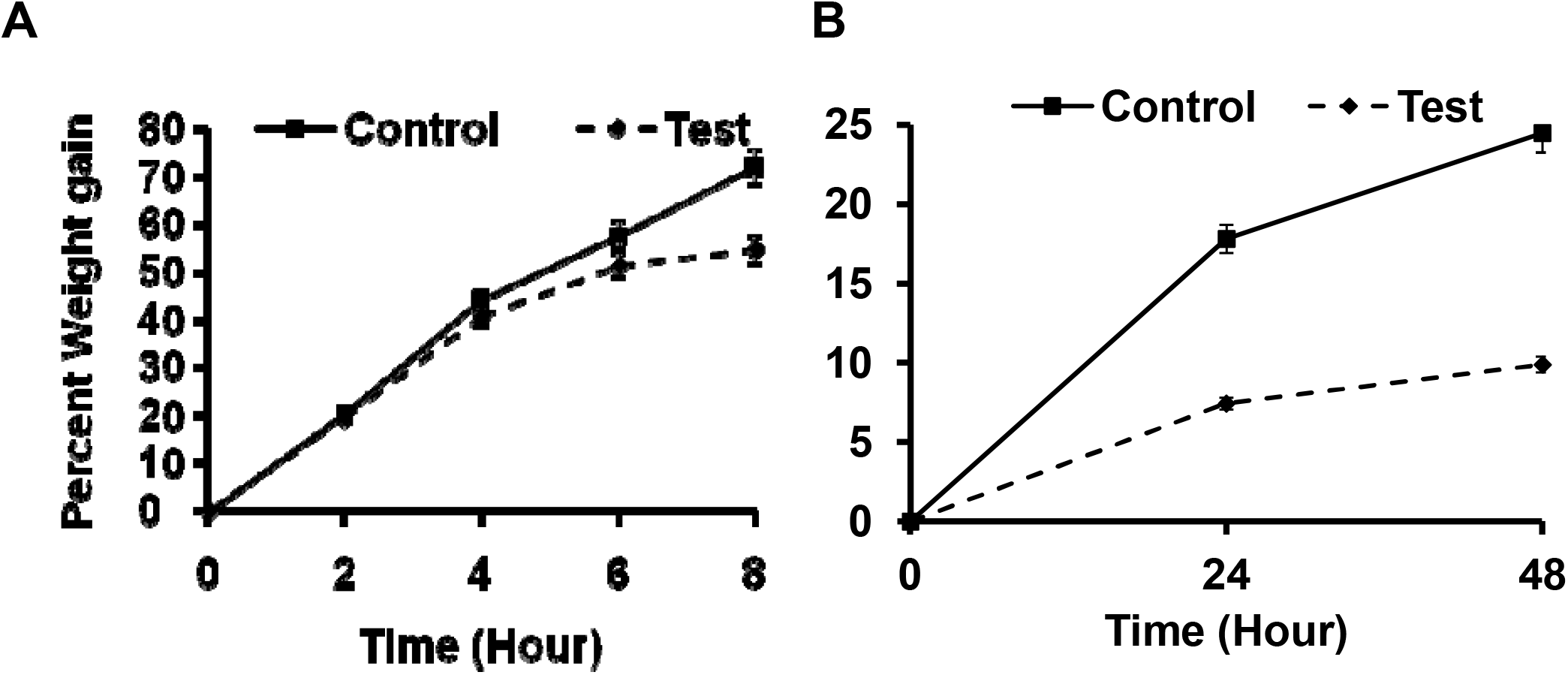
Effect of HarmGr9 blocking on host plant feeding. **(A)** Percent weight gain by the insects fed on tomato leaf disc coated with DEPC water (Control) and miglitol (Test) in 50 mM concentration. **(B)** Percent weight gain by the insects fed on tomato plant sprayed with DEPC water (Control) and miglitol (Test) in 50 mM concentration

In our second set up, the effect of HarmGr9 blocking on the feeding response, in the natural scenario, was checked by allowing larvae to feed on the entire tomato plant for 48 hours. For test insects, the plants that were provided were sprayed with 50 mM of miglitol solution while the control insects were fed on plants sprayed with DEPC water. The initial weight of all the insects was taken followed by recording the insect weight every 24 hours. The difference in PWG of the test and control insects was significantly high. After 48 hours of feeding, the total weight gain by the control insects was around 25% whereas for the test insects it was around 10% (Fig. 3B). These two experiments supported the earlier results and reconfirmed the potential of miglitol as a HarmGr9 blocker. Additionally, we observed that some larvae showed cannibalistic behavior when allowed to feed on the entire plant. It has been previously reported that the cannibalism in *H. armigera* is induced by various factors such as food availability, temperature and larval development stage (Tang et al., 2016). This cannibalistic tendency was only seen in test insects which allowed to fed on miglitol sprayed plant. As per our understanding, the reduced feeding response of larvae due to HarmGr9 blocking must have generated a false alarm situation of food scarcity for the insects which in turn induced cannibalism. This observation in future can be further investigated to better understand the behavioral responses of *H. armigera* to GR silencing. We consider that this particular response can be an additional advantage of GR blocking, apart from feeding inhibition, which can be explored in developing strategies for insect control.

### 3.4. Effect of RNAi mediated knockdown of HarmGr9 on insect feeding response

In order to strengthen our observations of HarmGr9 blocking and its effect on *H. armigera* feeding response, RNAi mediated knockdown of HarmGr9 was carried out. The larvae fed on HarmGr9-siRNA, followed by feeding assay and RNA extraction. The qRT-PCR analysis clearly showed a significant decrease in mRNA levels of HarmGr9 in test insects as compared to the control insects and indicated the successful knockdown of HarmGr9 (Fig. 4A). Post siRNA feeding, the feeding assay with agarose block, as mentioned in section 2.4, was carried out for 6 hours with test and control insects. The feeding response showed a significant inhibition in test insects as compared to control insects. The percent feed eaten by control insects was about 40% of the total feed whereas the test insects consumed only around 20% of the total feed (Fig. 4B). This observation is comparable to our previous results and thus reinforced our assertion of miglitol as a HarmGr9 blocker. In parallel to our previous results, siRNA fed insects also showed a significant reduction in PWG as compared to control insects (Fig. 4C).

**Figure 4:**
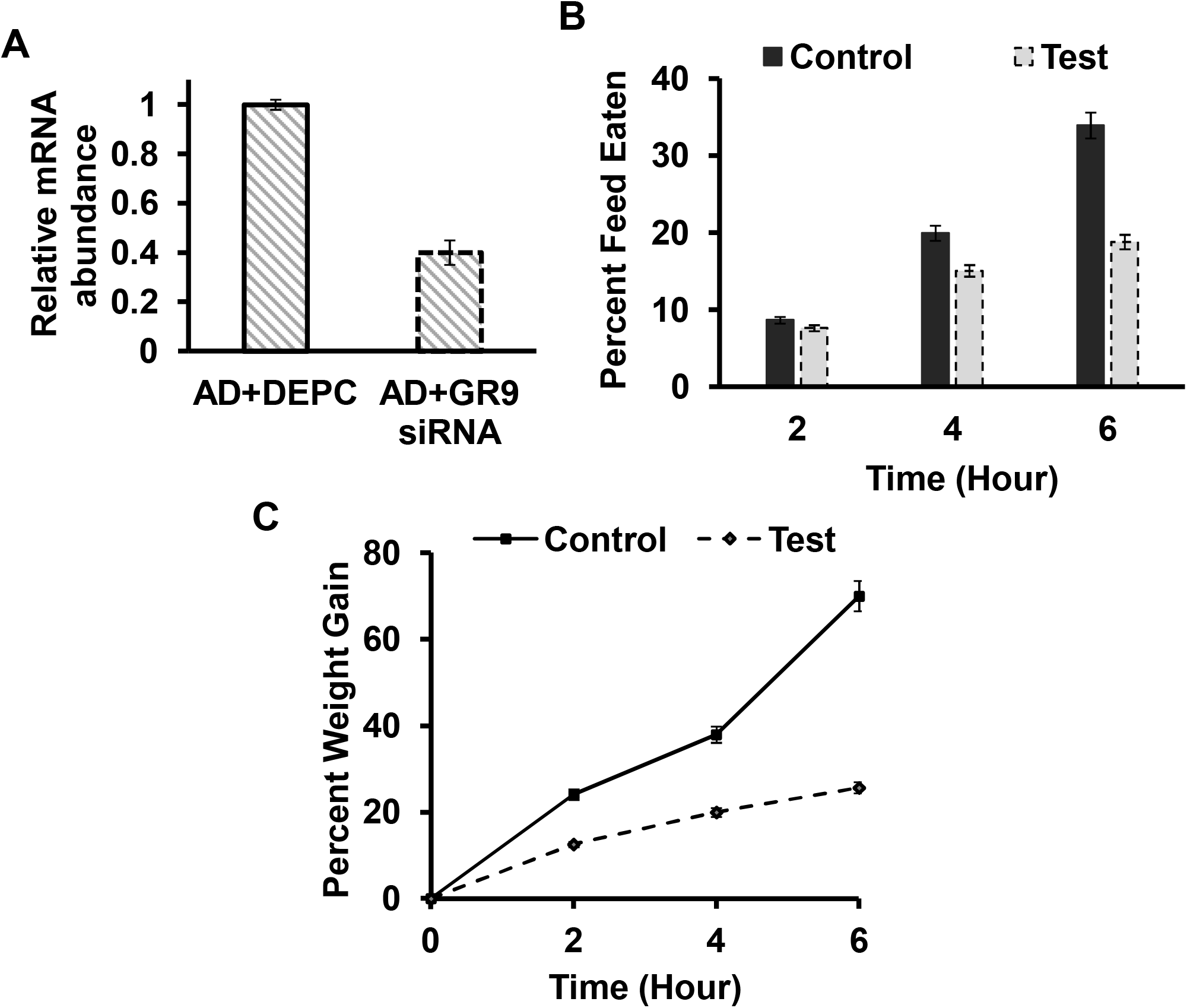
Effect of RNAi mediated knockdown of HarmGr9 on insect feeding response. **(A)** Relative abundance of HarmGr9 mRNA in control (AD + DEPC) and test (AD + HarmGR9-dsRNA) insects. **(B)** Percent feed eaten by test and control insects post dsRNA treatment. **(C)** Percent weight gain by the insects due to feeding post dsRNA treatment

Many of the previous studies that have used RNAi mediated knockdown of GRs in insects, as an approach to studying the GR functioning, have preferred injecting the siRNA in insects rather than feeding it to them (Utoguchi et al., 2011). We particularly used feeding approach to specifically target the mouth and foregut Gr9 in *H. armigera*. As per our understanding, tissue-specific blocking is essential for observing the miglitol mediated silencing of Gr9 and its corresponding effect on feeding responses of *H. armigera*. We believe, the effect of miglitol on Gr9 expressed in tissues other than mouth and foregut can be studied in future.

## 4. CONCLUSION

The GR is one of the essential chemosensory molecules responsible for governing various behavioral and physiological roles in *H. armigera*. HarmGr9 is one of such GRs that have a specific role in detecting the various host plant sugars and thus initiating the feeding behavior in *H. armigera*. In this article, we have followed a novel approach of silencing HarmGr9 using miglitol as a blocker and thus inhibiting the feeding potential of *H. armigera*. Based on the Ca^2+^ imaging study of the *in-vitro* expressed HarmGr9, it can be observed that miglitol has a significant potential of silencing the Gr9 activation. We have further explained this by analyzing the feeding response of larval insects exposed to miglitol. We propose the blocking of sugar-specific GRs in *H. armigera* by using imino sugar, as a novel prospective strategy for agricultural pest control.

## ACKNOWLEDGEMENT

The project work is supported by the research grant from the Department of Science and Technology - Science and Engineering Research Board (DST-SERB), Government of India under ECR/2015/000502 grant and UPE II grant, Savitribai Phule Pune University, Pune 411007, Maharashtra, India. The authors acknowledge Dr Alisha Anderson (CSIRO, Australia) for sharing the HarmGr9 clone and Dr Wei Xu (Murdoch University, Australia) for the technical help in planning cell-based experiments. Authors acknowledge Dr Tuli Dey from IBB, SPPU for the critical comments and editorial assistance.

## NOTE

The authors declare no competing financial interest.

